# Viability Discrimination of Bacterial Microbiomes in Home Kitchen Dish Sponges via Propidium Monoazide Treatment

**DOI:** 10.1101/2022.07.22.501210

**Authors:** Christina K. Carstens, Joelle K. Salazar, Shreela Sharma, Wenyaw Chan, Charles Darkoh

**Affiliations:** University of Texas Health Science Center, School of Public Health, Department of Epidemiology, Human Genetics and Environmental Sciences, Houston, TX, USA; Division of Food Processing Science and Technology, U.S. Food and Drug Administration, Bedford Park, IL, USA; University of Texas Health Science Center, School of Public Health, Department of Biostatistics and Data Science, Houston, TX, USA; University of Texas MD Anderson Cancer Center UTHealth Graduate School of Biomedical Sciences, Microbiology and Infectious Diseases Program, Houston, TX, USA

**Keywords:** Kitchen microbiome, Kitchen pathogens, Propidium monoazide treatment, Microbiome discrimination, Dish sponge pathogens

## Abstract

Dish sponges are known to support the survival and growth of human bacterial pathogens yet are commonly used by consumers to wash dishes and clean kitchen surfaces. Exposure to foodborne pathogens via sponge use may lead to foodborne illness, which is of particular concern among susceptible populations. Limitations associated with culture-independent or - dependent methods for bacterial community characterization have challenged the complete assessment of foodborne pathogen exposure risk presented by sponges. In this study, the bacterial microbiomes of five dish sponges were characterized to evaluate the presence of viable bacterial foodborne pathogens using propidium monoazide treatment, which is a novel approach in this medium. Total and viable sponge microbiomes were subsequently metataxonomically evaluated via targeted 16S rRNA sequencing. Select pathogen viability was confirmed using targeted selective enrichment. The comparison of total and viable sponge microbiome beta diversity indicated that sponge taxonomic abundance profiles did not vary significantly according to PMA treatment. The numbers of unique bacterial species (*p*-value = 0.0465) and unique bacterial foodborne pathogens (*p*-value = 0.0102) identified were significantly lower after PMA-treatment. A total of 20 unique bacterial foodborne pathogens were detected among sponge microbiomes regardless of PMA treatment. Three to six unique viable foodborne pathogens were identified in each sponge. *Escherichia coli* and *Staphylococcus aureus* were identified in all five viable sponge microbiomes evaluated. Viable *E. coli* were recovered from two of five sponges via targeted selective enrichment. These findings suggest that most sponge-associated bacterial communities may be viable and contain multiple viable bacterial foodborne pathogens.

**Importance:** Bacterial pathogens may exist in the domestic kitchen environment, threatening both sanitation levels and the health of residents. Dish sponges are commonly used kitchen tools that can harbor foodborne pathogens as they present adequate conditions for the survival and growth of bacteria. Using a contaminated sponge may lead to foodborne illness through direct contact with pathogens or via cross-contamination with food or other surfaces. Although bacterial foodborne pathogens have been identified in sponges, previous limitations in methodology have prevented the complete understanding of sponge microbiomes. This study used a novel application of a chemical reagent coupled with targeted amplicon sequencing to identify sponge communities and differentiate between viable and non-viable bacteria. Insights into dish sponge microbiomes and potential risks of bacterial foodborne pathogen exposure can inform food safety education programs to aid in the prevention of home-acquired foodborne illness and cross-contamination events.

## Introduction

Most foodborne illnesses in the United States are home-associated and sporadic [1]. Although most cases are self-limiting, severe outcomes are primarily caused by bacterial etiologies, including *Escherichia coli* O157:H7, *Salmonella enterica*, and *Listeria monocytogenes* [2]. Evidence indicates that low-income and racial-ethnic minority populations in the U.S. experience disproportionately higher rates of foodborne illnesses [3]. Studies also suggest that children from low-income families are at the most significant risk for numerous foodborne infections compared to those of higher income [4-6], which increases concern for foodborne pathogen presence, survival, and growth on home surfaces.

Bacteria are transported into the home via various routes and concentrate at the highest levels within kitchens. Bacterial contamination within kitchens can lead to cross-contamination of food, which is a leading cause of foodborne disease outbreaks [7]. The dominating bacterial presence in domestic kitchens is partially driven by the number of bacteria present on dish sponges, which harbor the highest coliform levels of home kitchen contamination sites [8]. As dish sponges can support bacterial survival and growth and transfer bacteria to other surfaces [9], they are considered bacterial reservoir disseminators. Bacterial pathogens, including *Salmonella* species, are commonly detected in used dish sponges [8, 10] and have been observed to survive within them for up to ten days. Despite this, one European study found that over 70% of consumers do not change their dish sponges until after three days of use or more [11]. Further, over 50% of consumers also use their dish sponges to clean the kitchen countertop, which may contribute to cross-contamination. Among a predominantly low-income population of parents in Texas, 71% of respondents also reported using a reusable cleaning tool (i.e., dish sponges and cloths) to clean the kitchen counter after food preparation [12].

Sponge storage and treatment conditions by consumers play a critical role in bacterial survival and growth. Evidence indicates that dry storage of dish sponges may reduce the risk of cross-contamination as low sponge moisture levels have been linked to decreased bacterial survival over time [11]. Treatment of sponges such as microwaving or dishwashing has been observed to significantly reduce the number of aerobic bacteria present [13], and microwaving has also been observed to reduce bacterial community diversity [14]. However, other research has found that regular sponge sanitization has no significant effect on the number of bacteria present [15], and sponge cleaning methods have been reported as more effective when conducted in the laboratory than in the home environment [10]. Consumer efforts to sanitize dish sponges may be inhibited by the high levels of organic matter and food residues that build up after frequent and extended use. Sponge sanitization has also been linked to significant differences in microbial community composition [14] and an increase in the presence of bacteria linked to the production of unpleasant odor [15]. Sponge treatment with a 4000 ppm hypochlorite solution overnight (16-20 h) has been demonstrated to reduce total bacteria and *Salmonella* spp. levels and prevent bacterial regrowth [11]. Consumer attempts to sanitize dish sponges may contribute to the proliferation of bacteria that are resistant to sanitization, including those with malodorous or pathogenic properties, depending on the cleaning method used.

Although consumer use of dish sponges can lead to bacterial reservoir generation and cross-contamination, the full scope of foodborne pathogen exposure risk from sponges is not understood. For example, studies that have relied on culture-based methods to detect pathogens within sponges have been limited by the choice of specific pathogen targets and their culturability. Further, studies that evaluated sponge microbial communities via culture-independent methods (i.e., genetic sequencing) [14-16] have been unable to determine the risk of pathogen exposure as species level identifications were not obtained and microbial community viability was not assessed, despite the availability of chemical-based viability discrimination methods. As metataxonomic sequencing cannot indicate if sequenced DNA originated from live or dead bacteria, viability discrimination is a process by which DNA from solely intact, viable bacterial cells is isolated before downstream genetic sequencing. Several studies of the microbial communities present within the indoor built environment, such as cleanrooms and public transit systems [17, 18], have utilized propidium monoazide (PMA) to differentiate between DNA from viable and non-viable cells (relic DNA). PMA is a DNA-binding dye that can penetrate compromised cell membranes but is excluded from viable Gram-positive and Gram-negative cells [19]. Upon photoactivation, PMA covalently binds to available relic DNA, renders it insoluble, and allows for selective downstream gene amplification and targeted 16S rRNA sequencing of DNA from exclusively viable cells.

Viability discrimination via PMA treatment paired with subsequent targeted 16S rRNA sequencing is a novel approach to evaluate dish sponge bacterial communities and detect viable bacterial foodborne pathogens. This study aimed to examine the total and viable bacterial microbiomes of dish sponges to assess viable foodborne pathogen presence. This research aimed to 1) estimate the taxonomic diversity and distribution of the total and viable dish sponge bacterial communities and 2) differentiate viable and non-viable foodborne pathogens present on the dish sponges via culture-independent and culture-dependent methods.

## Methods

### PMA Optimization

Bacterial strains were obtained from the U.S. Food and Drug Administration (FDA) strain stock inventory (Bedford Park, IL). *Listeria monocytogenes* strain Scott A [20] and *E. coli* strain K12 (ATCC 25253) working stocks were maintained on tryptic soy agar (TSA; BD Difco, Franklin Lakes, NJ). Cultures were grown in tryptic soy broth (TSB; BD Difco) incubated at 37 °C for 16-18 h.

DNA extraction was conducted using the DNeasy Blood & Tissue Kit (Qiagen Inc., Germantown, MA). Extracted DNA was quantified using the Qubit dsDNA HS Assay Kit (Invitrogen, Carlsbad, CA). DNA extraction and quantification were performed according to the manufacturer’s instructions with a 100 uL starting volume. All genomic DNA was stored at - 20°C until further analysis.

Propidium monoazide (PMA; Biotium, Inc., Fremont, CA) treatment concentrations of 30 and 50 μM and light exposure times of 60, 90, 120, 150, and 180 s were tested using *L. monocytogenes* genomic DNA. All reactions were conducted in replicate (n = 40) and consisted of 1 ng/μL template DNA, nuclease-free water, and either 0, 30, or 50 μM PMA in a reaction volume of 100 μL. Reaction tubes were covered with aluminum foil and incubated at ambient temperature for 5 min with inversion at 1 min increments. Samples were then transferred to the reaction surface, which consisted of aluminum foil placed over ice. Light exposure treatment was conducted using a 600 W halogen lamp at least 20 cm away from the reaction tubes. The reaction surface was shaken at 30 s increments during light exposure to ensure reaction homogeneity. After treatment, samples were placed on ice for 5 min. The temperature of the reaction surface was tracked using a digital temperature probe (Fisherbrand, Waltham, MA). PMA addition and reactions were conducted in a dark room.

qPCR targeting the *L. monocytogenes actA* gene was performed on a 7500 Fast Real-Time PCR system (Applied Biosystems, Foster City, CA). Reactions were conducted in triplicate using PowerUp SYBR Green Master Mix (Thermo Fisher Scientific, Inc, Waltham, MA) with 1 μL of 10 nM primer *actA*-F (5’-GATTTATGCGTGCGATGA-3’), 1 μL of 10 nM primer *actA*-R (5’-TTACCTCGCTTGGTTGCTC-3’), and 8 μL DNA template in a total volume of 20 μL. Cycling parameters were: 50°C for 2 min, 95°C for 2 min, 40 cycles of 95°C for 15 s, 56°C for 30 s, and them 72°C for 1 min. A negative control that consisted of all reagents aside from the DNA template was included in each run. Two independent trials of PMA optimization experiments, including DNA extraction, PMA treatment, and qPCR, were conducted.

### Sponge Acquisition, Processing, and Biomass Quantification

Sponge sample collection and processing were conducted as described previously [21] from the homes of predominantly low-income families with at least one child enrolled in a Houston Independent School District elementary school. Parents were recruited via the distribution of flyers in partnership with the nonprofit nutrition intervention Brighter Bites, which operates in schools were at least 75% of the students are receiving free or reduced-price lunch [22]. After sponge sample collection, sponges were homogenized by hand with 50 mL 1× phosphate-buffered saline for 1 min. Sample collection was approved by the University of Texas Health Science Center Institutional Review Board Committees for Protection of Human Subjects. Bacterial genomic DNA was extracted from five sponges as described above. Estimation of the total number of 16S rRNA gene copies present in each sample was conducted via qPCR as previously described targeting the 16S rRNA gene. Modifications to reaction volumes included 0.5 μL of 10 μM forward primer 26F2a, 0.5 μL of 10 μM reverse primer 534R2 [23], and 4 μL of DNA template. Cycling conditions were 50°C for 2 min, 95°C for 2 min, 40 cycles of 95°C for 15 s, 56°C for 30 sec, and then 72°C for 2 min. DNA was extracted from *E. coli* K12, quantified, serially diluted 1:10, and enumerated via plate count assay to generate the standard curve.

### Sponge Treatment

Viability discrimination was conducted on sponge samples from each household via PMA treatment. PMA reactions were performed for each sponge in replicate and consisted of 50 μM PMA with a light exposure time of 180 s, as previously described. Replicate PMA-free samples for each sponge also underwent PMA reaction conditions. DNA extraction and quantification from each sample (n = 20) were conducted immediately after PMA treatment.

### Amplification of 16S rRNA Genes, Library Construction, and Sequencing

DNA extracted from each sponge sample (n = 20) was used as template DNA for amplification of bacterial 16S rRNA genes. Primer pairs to target the V1-V3 region were randomly distributed equally amongst the DNA samples as described in Salazar et al. [23]. PCR products were visualized via agarose gel electrophoresis, quantified, and purified using AMPure XP beads (Beckman-Coulter, Indianapolis, IN) according to the manufacturer’s instructions. The Nextera XT Kit (Illumina, San Diego, CA) was used to index the 16S rRNA PCR products as previously described [23]. Indexed PCR products were normalized to 2 nM, pooled, and diluted to 12 pM. The library was spiked with 10% of 12.5 pM PhiX and sequenced using 150 cycles of MiSeq version 3 chemistry (Illumina).

### Targeted Enrichment and Colony Identification

Any samples for which *E. coli* or *Salmonella* spp. were identified via 16S rRNA sequencing were enriched according to the U.S. FDA Bacteriological Analytical Manual [24] modified with a sample volume of 100 μL in each primary enrichment. Presumptive colonies of *E. coli* or *Salmonella* were streaked onto TSA and verified via 16S rRNA sequencing as previously described using colony PCR to amplify the 16S rRNA gene fragment.

### Data Analysis

Average ΔC_*T*_ values (PMA-free – PMA-treated samples) and standard deviations were calculated across independent trials for the PMA optimization experiments. The linear regression line equation obtained from the comparison of C_t_ values to log colony-forming units (CFU) associated with the *E. coli* culture was used to compute the biomass of PMA-free bacterial communities for each sponge.

Analysis of the 16S rRNA sequencing of sponge samples was conducted as previously described [23]. Briefly, raw, paired-end reads were merged, then filtered based on quality (Q30, 99.9% accuracy) and length (minimum 300 base pairs). Sequences were aligned to the Ribosomal Database Project (RDP) [25] and the National Center for Biotechnology Information (NCBI) 16S databases [26] and taxonomically classified at the species level using Kracken2 [27]. Bracken [28] was used to estimate the relative abundance of each bacterial taxa. A filter of 0.05% relative abundance was used to exclude sequencing artifacts and singletons. Any bacterium known to cause foodborne illness [29] was considered a bacterial foodborne pathogen. As identified *E. coli* species were of unknown serotype, *E. coli* species were included as possible foodborne pathogens when identified. Relative abundance values were averaged across replicate samples to summarize the number of unique bacterial species and bacterial foodborne pathogens identified among sponge microbiomes according to PMA-treatment. The median numbers of unique bacterial species and bacterial foodborne pathogens identified within PMA-free and non-PMA treated sponge samples were statistically compared via the Mann-Whitney U test. Averaged relative abundance values for each taxon were also used to compute changes in relative abundance between PMA-free and PMA-treated bacterial communities for each sponge.

The “vegan” package for R [30] was used to calculate alpha and beta diversity parameters for each replicate sponge sample. Observed OTU counts, the Chao 1 index, Shannon’s diversity index, Simpson’s index, and Simpson’s reciprocal index were computed to examine sample alpha diversity and summarized across replicates with averages and standard deviations. The beta diversity of sponge microbiomes was visually compared with non-metric multidimensional scaling (NMDS) using Bray-Curtis dissimilarity. A permutational multivariate analysis of variance (PERMANOVA) test was used to evaluate beta diversity across PMA-free and PMA-treated sponge samples using the “vegan” package. A *p*-value of <0.05 was considered statistically significant for all statistical tests. Data analysis was conducted using the R language [31] (version 3.6.3, R Foundation for Statistical Computing, Vienna, Austria) and RStudio software [32] (version 4.1.2; RStudio, Inc., Boston, MA). Data was also visualized using Tableau software (version 2020.2.4, Tableau Software, Inc., Seattle, WA) [33].

### Data Availability

The sequence files and metadata generated in this study has been deposited to NCBI under Bioproject PRJNA83087, Biosamples SAMN28037372-SAMN28037381.

## Results

Propidium monoazide (PMA) treatment was evaluated on genomic DNA extracted from *L. monocytogenes* Scott A to optimize PMA reactions for bacterial community viability discrimination before use with dish sponge samples (Supplemental Figure 1, Additional File 1). The large differences in cycle thresholds (C_t_ values) observed between equivalent PMA-treated and PMA-free samples across concentrations and reaction times evaluated indicate that PMA was able to inactivate the amplification of DNA from heat-treated *L. monocytogenes* regardless of concentration or time. As PMA efficiency did not vary according to the concentration or reaction times assessed, a concentration of 50 μM PMA and a reaction time of 180 s were chosen to ensure sufficient PMA availability and time for DNA binding.

The biomass of the total bacterial community within each sponge was estimated using qPCR targeting the 16S rRNA gene and comparison with a standard curve (Supplemental Figure 2, Additional File 1). Sponge biomass levels ranged from 0.25 to 5.18 log CFU. The biomass level for one sponge was low (0.25 log CFU), while the biomass for the remaining four sponges was at least 1.73 log CFU higher (1.98 to 5.18 log CFU). When dish sponges were collected from each household, observations were made regarding the state of each sponge. Notably, high-biomass sponges were visibly wet when collected, whereas the low-biomass sponge was dry.

### Sponge Microbial Diversity

A total of five dish sponge bacterial communities were subjected to viability discrimination via PMA treatment. Alpha diversity parameters were computed for total (PMA-free) and viable (PMA-treated) sponge bacterial communities (Supplemental Table 1, Additional File 1). The alpha diversity of sponge microbiomes, as measured by average Shannon’s diversity index across replicates, ranged from 1.29 to 2.36 for total sponge microbiomes and 1.24 to 2.99 for viable sponge microbiomes. The average number of operational taxonomic units (OTU) observed ranged from 16.0 to 29.0 for total sponge microbiomes and 8.0 to 37.0 for viable sponge microbiomes. Differences between total and viable bacterial communities were evaluated via non-metric multidimensional scaling (NMDS) using Bray-Curtis dissimilarity (Figure 1). However, the taxonomic composition of sponge microbiomes was not observed to vary significantly according to PMA treatment.

**Figure 1.**
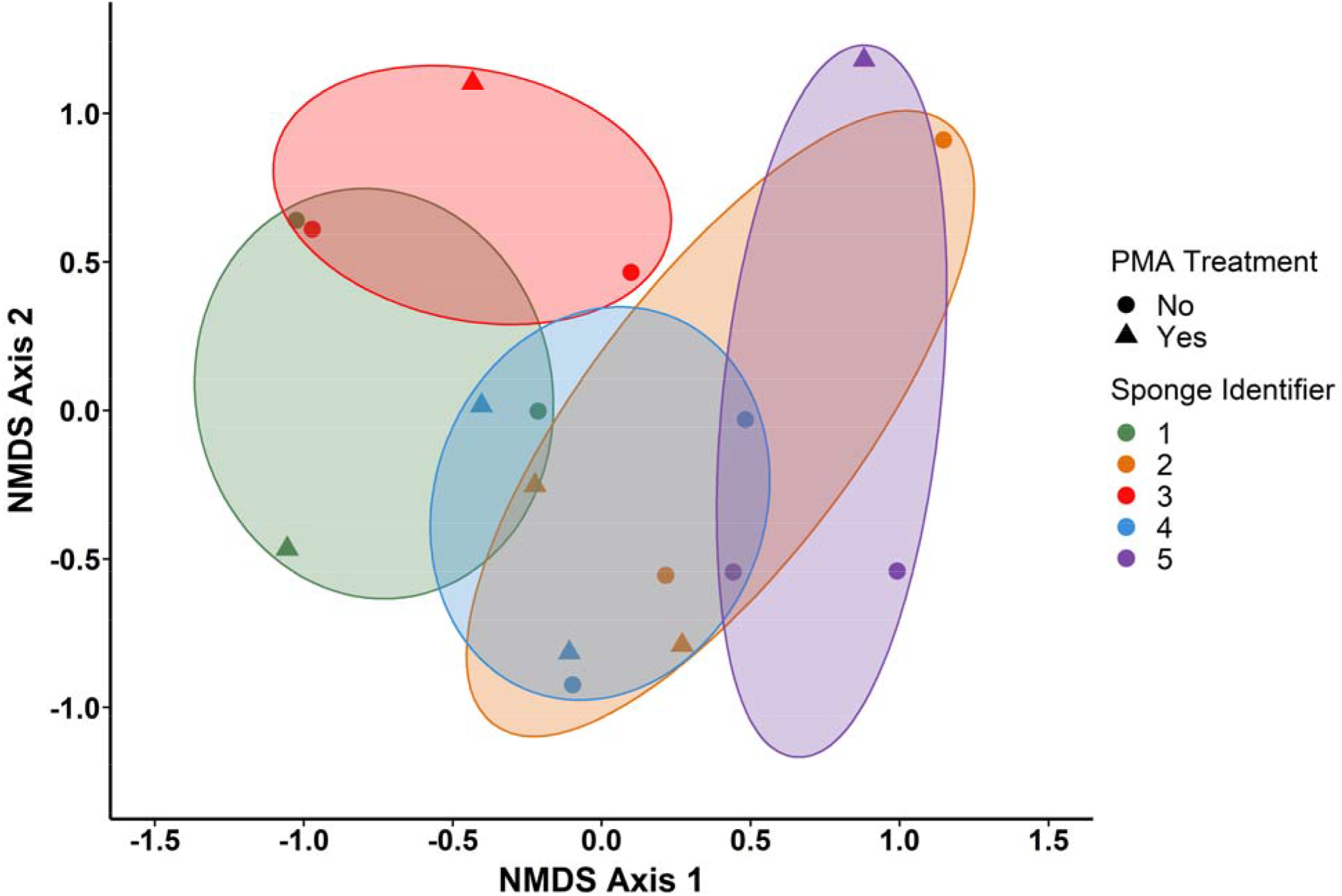
Non-metric multidimensional scaling (NMDS) of Bray-Curtis dissimilarity for sponge bacterial communities according to propidium monoazide (PMA) treatment. Beta diversity computations were conducted for replicate sponge samples using the relative abundances for each taxon.

### Sponge Microbiome Response to PMA-Treatment

The relative abundances of bacterial species identified in total and viable sponge microbiomes were averaged across replicate samples and compared to PMA-treatment. A total of 142 unique bacterial species were identified among total sponge microbiomes, and the number of species per total sponge bacterial community ranged from 34 to 49 (median: 44.0). Of the 142 unique species identified, 103 only appeared in one out of the five total sponge microbiomes evaluated. The most common species identified were *E. coli* and *Klebsiella pneumoniae* (5 out of 5 sponges each). Among viable sponge microbiomes, a total of 97 bacterial species were identified, and the number of species ranged from 11 to 40 (median: 28.0). *E. coli* and *Staphylococcus aureus* were the most common species identified among viable sponge bacterial communities (5 out of 5 sponges each). Most species identified among viable sponge microbiomes also only appeared once across the five communities evaluated (74 species). A significantly higher number of unique bacterial species was identified among total sponge microbiomes (median: 44.0 species) compared to viable sponge microbiomes (median: 28.0 species; *p*-value = 0.0465).

Dominant species in each sponge microbiome were evaluated according to their mean relative abundance across replicate samples and PMA-treatment status. Among total sponge bacterial communities, *K. pneumoniae* was the dominant species in two out of the five sponges evaluated, with mean relative abundances of 29.6% (sponge 4) and 39.7% (sponge 5). Each of the five total sponge microbiomes contained *K. pneumoniae* with a mean relative abundance ≥13%. According to the highest mean relative abundance, the dominant species in the remaining three total sponge bacterial communities were *Brucella suis* (sponge 1; 17.2%), *Acinetobacter baumannii* (sponge 2; 61.3%), and *Methylobacterium* spp. 17Sr1-28 (sponge 3; 50.1%). In viable sponge microbiomes, *K. pneumoniae* was also the dominant species in two instances, with mean relative abundances of 29.1% (sponge 3) and 46.5% (sponge 4); however, it was only present in three of the five communities after PMA treatment. *Streptomyces* spp. ICC4 (sponge 1; 15.8%), *Acinetobacter baumannii* (sponge 2; 53.6%), and *Ralstonia pickettii* (sponge 5; 32.3%) were the dominant species in the remaining three viable sponge microbiomes according to mean relative abundance levels.

The species with the top ten largest absolute changes in relative abundance between total and viable sponge microbiomes were examined to assess the species most affected by PMA-treatment (Figure 2). Among species with the top ten largest absolute changes across sponges, 20 unique species decreased in relative abundance after PMA-treatment, while 17 increased in relative abundance. *K. pneumoniae* (3 sponges) was the most common species to decrease in relative abundance after PMA-treatment, while *S. aureus* (5 sponges) and *Streptomyces* spp. ICC4 (3 sponges) most frequently increased in relative abundance. The relative abundance of *E. coli* increased in three sponge bacterial communities after PMA-treatment and decreased in two communities.

**Figure 2.**
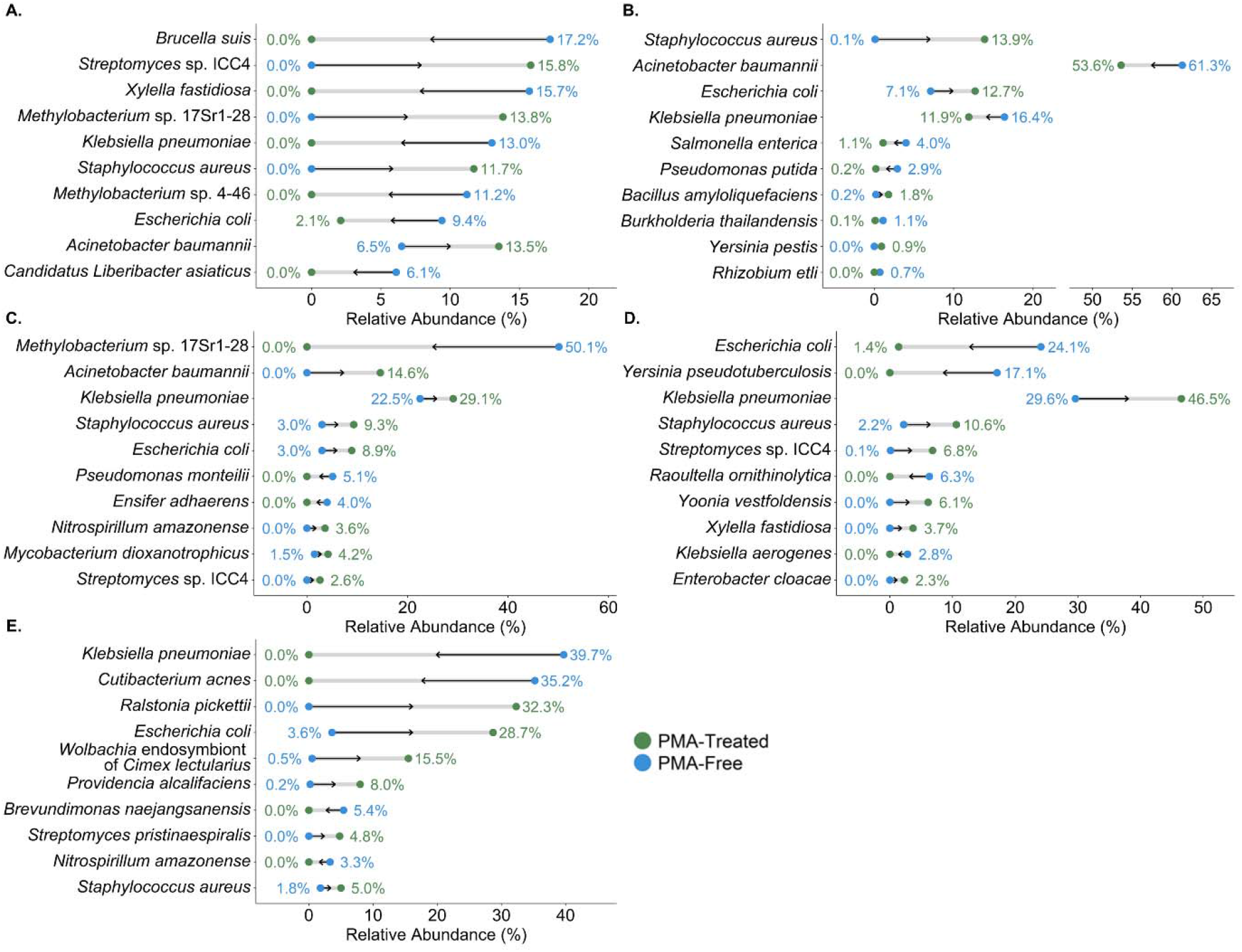
Taxa with the top ten largest changes in relative abundance between propidium monoazide (PMA) free and PMA-treated samples for sponge 1 (A), sponge 2 (B), sponge 3 (C), sponge 4 (D), and sponge 5 (E). Changes in relative abundance between PMA-free and PMA-treated samples were computed using the averaged relative abundances of each taxon across replicate samples.

Species with <1.0% changes in relative abundance after PMA-treatment were identified. Most of these species were present in small relative quantities within the total sponge bacterial communities. A total of 124 unique species were identified with at least one instance of a <1.0% change in relative abundance across the five sponge microbiomes evaluated. The number of unique species per sponge with small changes in relative abundance after PMA-treatment ranged from 26 to 36 (median: 35.0). Species that frequently experienced these small changes in relative abundance included *Alkalitalea saponilacus, Betaproteobacteria bacterium* GR16-43, *Draconibacterium orientale, Pasteurella multocida, Raoultella ornithinolytica*, and *Thermoanaerobacterium thermosaccharolyticum* (3 sponges each, respectively). The most common bacterial foodborne pathogens with small changes in relative abundance included *Bacillus thuringiensis* (3 sponges), *Enterobacter cloacae* (3 sponges), *Bacillus* spp. FJAT-45348 (2 sponges), *Providencia alcalifaciens* (2 sponges), and *Streptococcus pneumoniae* (2 sponges).

Species identified in the PMA-free sponge bacterial communities but not in the respective PMA-treated bacterial communities were examined to ascertain species that were not likely viable in the total sponge microbiome. Across the five sponges evaluated, a total of 136 unique species were identified in a total sponge microbiome and not in the respective viable sponge microbiome in at least one instance. The number of species identified in the total community that was not identified in the respective viable community per sponge ranged from 19 to 40 (median: 38.0). The most common species that were identified as likely non-viable in sponge microbiomes via PMA treatment included *R. ornithinolytica* (4 sponges), *B. bacterium* GR16-43 (3 sponges), and *Nitrospirillum amazonense* (3 sponges). *B. thuringiensis* (3 sponges) was the most frequently identified bacterial foodborne pathogen in total sponge microbiomes, which was subsequently not identified in viable sponge microbiomes.

### Foodborne Pathogen Presence and Response to PMA-Treatment

A total of 20 unique bacterial foodborne pathogens were identified across the five sponge bacterial communities, regardless of PMA-treatment. Among total sponge microbiomes, 19 unique foodborne pathogens were identified, ranging from 7 to 13 per sponge (median: 8.0). *K. pneumoniae* was the dominant foodborne pathogen in four total sponge bacterial communities (sponges 2-5) with relative abundance >16%, while *B. suis* was the dominant foodborne pathogen in the total microbiome of sponge 1. A total of eight unique bacterial foodborne pathogens were identified among viable sponge microbiomes; at least three were identified within each dish sponge evaluated, with a maximum of six (median: 5.0). The dominant foodborne pathogens identified among viable sponge microbiomes according to the mean relative abundance included *S. aureus* in sponges 1 and 2 (11.7 and 13.9%, respectively), *K. pneumoniae* in sponges 3 and 4 (29.1 and 46.5%, respectively), and *E. coli* in sponge 5 (28.7%). The number of unique foodborne pathogens identified per sponge was significantly higher among total sponge microbiomes (median: 8.0 species) than viable sponge microbiomes (median: 5.0 species; *p*-value = 0.0102).

The most common foodborne pathogens identified among total sponge bacterial communities included *E. coli* (5 sponges), *K. pneumoniae* (5 sponges), *E. cloacae* (4 sponges), *P. alcalifaciens* (4 sponges), and *S. aureus* (4 sponges). Nine unique foodborne pathogens were only identified in one out of the five total sponge microbiomes evaluated. *E. coli* (5 sponges), *S. aureus* (5 sponges), *E. cloacae* (3 sponges), *K. pneumoniae* (3 sponges), and *P. alcalifaciens* (3 sponges) were the most common foodborne pathogens identified among viable sponge bacterial communities. *Streptococcus pyogenes* was only identified in one of the five viable sponge microbiomes evaluated (sponge 4). Viable *E. coli* was identified in all five sponge bacterial communities and was recovered via targeted selective enrichment from sponges 1 and 5. *S. enterica* was identified in the viable bacterial communities for sponges 1 and 2 via PMA-based viability discrimination with relative abundance <3.0% but was not recovered via targeted selective enrichment.

Changes in relative abundance for the 20 identified foodborne pathogens within sponge bacterial communities after PMA-treatment are displayed in Figure 3. Across the five sponge microbiomes, a total of 18 unique foodborne pathogens decreased in relative abundance after PMA treatment in at least one instance. In contrast, eight unique foodborne pathogens increased in relative abundance in at least one instance. The foodborne pathogens that most frequently decreased in relative abundance were *B. thuringiensis* and *K. pneumoniae* (3 sponges, respectively). Conversely, *S. aureus* (5 sponges), *E. cloacae* (3 sponges), *E. coli* (3 sponges), and *P. alcalifaciens* (3 sponges) most frequently increased in relative abundance after PMA treatment.

**Figure 3.**
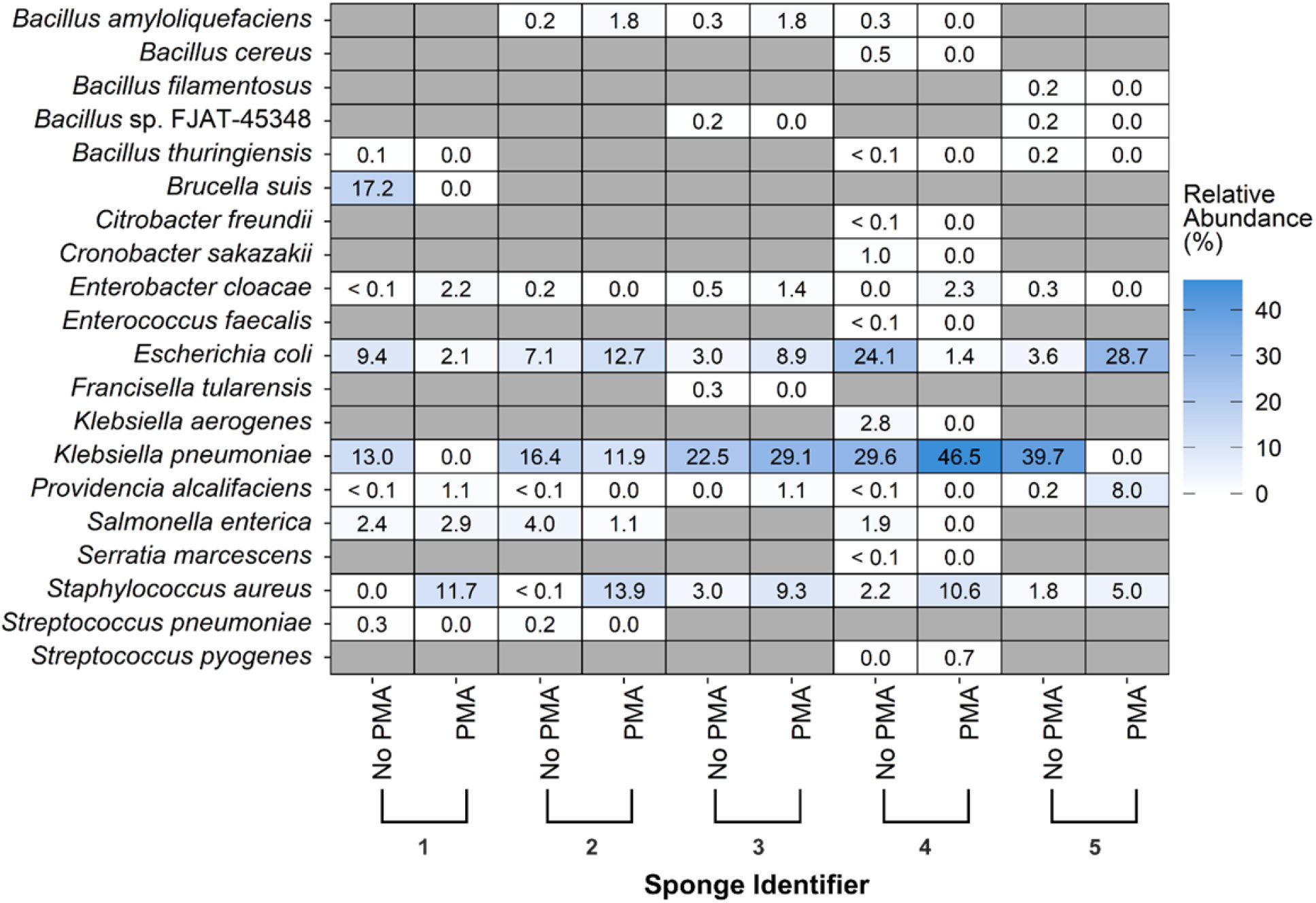
Changes in relative abundance for identified bacterial foodborne pathogen species between propidium monoazide (PMA) free and PMA-treated sponge samples. Numerical labels on the x-axis correspond to sponge identifiers (1-5) and PMA treatment status (PMA-free or PMA-treated). Gray tiles indicate the respective pathogen was not identified in either the PMA-free or PMA-treated sample for the corresponding sponge. Changes in relative abundance between PMA-free and PMA-treated samples were computed using the averaged relative abundances of each taxon across replicate samples.

## Discussion

The total and viable bacterial communities of five dish sponges were evaluated using 16S rRNA sequencing paired with viability discrimination via propidium monoazide (PMA) treatment and targeted selective enrichment. The dish sponges assessed in this study were a subset of environmental samples collected from the households of ten predominantly low-income families, as reported elsewhere [21]. Although past studies have used culture-dependent or independent methods to examine the bacteria in dish sponges, this is the first study to evaluate dish sponge microbiomes using PMA-based viability discrimination and targeted selective enrichment. The approach used in this study for viability discrimination via PMA treatment avoided traditional limitations associated with culture-based methods, including bacterial target selection and culturability. In addition, advancements in 16S rRNA databases enabled the identification of dish sponge microbiomes to the species level.

A wide range of biomass levels was observed across the five sponges evaluated in this study. The estimated number of bacterial cells per sponge ranged from 0.25 to 5.18 log CFU, although four of the five sponges had ≥1.98 log CFU. Studies that have used culture-based methods to enumerate bacteria in dish sponges obtained from domestic environments have found average quantities of ∼6-7 log CFU per sponge [34, 35]. Heterotrophic bacteria counts from kitchen sponges have also been observed to range from ∼2-8 log CFU [35], similar to the range observed in the present study. The variation in sponge biomass levels observed in the present study may be due to various factors, such as the frequency or duration of sponge use or sponge cleaning treatments. Consumers have reported microwave heating and rinsing with hot water and soap to clean sponges [15]; however, this study did not capture information on cleaning procedures or sponge use history. Although specific treatment and usage details were not obtained, observations of sponge moisture levels were made during sponge collection. Notably, the one sponge that appeared dry when collected contained the lowest biomass level identified. Previous research of artificially contaminated dish sponges with various water absorption properties indicated that the survival of *Salmonella* spp. over time was greater in sponges with slower drying times [11]. Dry storage of dish sponges may reduce bacterial loads; however, the addition of water and potential nutrients at the point of reuse may lead to rapid sponge recolonization by surviving bacteria.

According to mean Shannon’s diversity index, the alpha diversity for total sponge microbiomes ranged from 1.29 to 2.36, and mean observed OTU counts ranged from 16.0 to 29.0 per sponge. Other studies that used culture-independent methods to examine used dish sponges acquired from residential homes found a higher range of observed OTUs [14] and a wider range of Shannon diversity index values (0.93-5.07) [15] than those observed in the present study. Aside from methodological differences (shotgun sequencing and 454-pyrosequencing of 16S rRNA, respectively), these differences may be attributable to the lower number of sponges evaluated in this study, which may have limited the range of sponge microbiome diversity observed. Regarding sponge beta diversity, the taxonomic abundance profiles of total and viable sponge microbiomes examined in this study were not observed to vary significantly. However, the median numbers of unique bacterial species and bacterial foodborne pathogens identified within viable sponge microbiomes were significantly lower than those observed in total sponge microbiomes. Although the numbers of species varied, only ten species across all five sponges examined experienced at least one change in relative abundance ≥15% after PMA-treatment. This lack of variation in bacterial community composition may indicate that total and viable sponge microbiomes have similar taxonomic abundance profiles. Cellular features observed via fluorescence *in situ* hybridization analysis have demonstrated that most bacterial cells present in used dish sponges are in an active state of growth [15]. As dish sponges can provide conditions necessary for bacterial survival and proliferation, including moisture and nutrients, a high degree of bacterial viability within dish sponges is likely.

Common species identified across all total sponge microbiomes included *E. coli* and *K. pneumoniae*. Two previous studies of the microbiomes of domestically used dish sponges conducted in Germany did not find that members of the *Enterobacteriaceae* family were dominant in dish sponge bacterial communities. Specifically, the most frequently identified genera in one study were *Acinetobacter, Enhydrobacter, Agrobacterium, Pseudomonas*, and *Chryseobacterium*, while in the other, *Acinetobacter, Moraxella*, and *Chryseobacterium* were most frequently observed [15]. A study of kitchen surface microbiomes in the United States (Boulder, Colorado) similarly found that members of the family *Moraxellaceae* (40.4%) were dominant in the one sponge evaluated [16]. However, the U.S. study also found that the bacterial family with the second-highest relative abundance in the sponge microbiome was *Enterobacteriaceae* (16.8%). Several factors may contribute to the differences in the composition of dish sponge bacterial communities observed in this study. Food and water are two major sources of bacteria in kitchen microbiomes [16]. Therefore, variations in food preferences and water supplies across geographic locations could contribute to the observed differences in taxonomic abundance. Other external factors such as human residents or hygiene behaviors, and internal factors such as sponge size and material, may also contribute to these differences.

Bacterial genera that include foodborne pathogens identified among total sponge microbiomes in the present study have also been observed in other studies with low relative abundance (≤2%), including *Brucella, Citrobacter, Enterobacter, Escherichia, Klebsiella, Salmonella, Staphylococcus*, and *Streptococcus* [14, 15]. In the present study, *Brucella suis, E. coli, Klebsiella aerogenes, K. pneumoniae, S. enterica*, and *S. aureus* were identified among total sponge microbiomes with at least one instance of a relative abundance >2%. Among viable sponge microbiomes, *E. cloacae, E. coli, K. pneumoniae, Providencia alcalifaciens, S. enterica*, and *S. aureus* were identified with at least one instance of a relative abundance >2%. Metataxonomic sequencing of dish sponges artificially inoculated with a food soil suspension (i.e., 0.1% poultry soil, 0.1% egg-based soil, 1% lettuce soil) and kitchen-isolated bacteria (including *Staphylococcus, Salmonella*, and *Campylobacter* species) demonstrated that pathogens remained minority contributors to sponge bacterial communities over seven days of storage at room temperature [11]. Although foodborne pathogens have been observed as minor components of both naturally occurring and artificially inoculated dish sponge communities, the same pattern was not observed in this study. The food safety habits of dish sponge users may play a role in sponge contamination levels. A lack of serious concern for food contamination with germs among consumers has been linked to an increased variety of foodborne pathogens within home kitchens [21]. Competition dynamics of bacterial communities, as well as varying bacterial sources, initial loads, nutrient availability, and storage conditions, may also have led to the more dominant contributions of foodborne pathogens in the total and viable microbiomes observed in this study.

Viability discrimination using PMA treatment revealed that each dish sponge evaluated contained at least three unique bacterial foodborne pathogens. Viable foodborne pathogens identified in this study, including *E. cloacae [36], E. coli* [8, 10, 35, 37], *Klebsiella* spp. [36, 37], *Salmonella* spp. [10], and *Staphylococcus* spp. [8, 35, 36] have also been recovered from used dish sponges via culture-based methods. *Salmonella* spp. has even been observed to proliferate during the first day after artificial inoculation onto dish sponges stored at room temperature [11]. Although viable *E. coli* (5 sponges) and *S. enterica* (2 sponges) were identified using PMA-based viability discrimination, only *E. coli* was recovered from two sponges via targeted selective enrichment. The traditional limitations of culture-based methods, such as the inability to detect microorganisms that are viable but not culturable, under environmental stress, or out-competed in the bacterial community, may have inhibited the recovery of these bacterial targets.

In this study, dish sponges were observed to harbor viable bacterial foodborne pathogens; therefore, using sponges may present a direct exposure risk to these pathogens. Dish sponges are commonly used for cleaning purposes aside from washing dishes [11]. Contamination of dish sponges and cloths with *S. aureus* is also significantly associated with the same type of contamination on kitchen counters, sinks, refrigerator shelves, and refrigerator handle within the same kitchen [8]. Consequently, contaminated dish sponges may also serve as vehicles for cross-contamination of food and other kitchen surfaces. Consumers should be educated on the risks of bacterial foodborne pathogen exposure and the potential for cross-contamination linked to dish sponge use. Eliminating sponge use for dishwashing and cleaning would prevent the associated risks of direct pathogen exposure and cross-contamination; however, this preventive measure may not be economically feasible for low-income communities. Handwashing before and after handling sponges may reduce the risks of direct pathogen exposure. Frequent dish sponge replacement may also mitigate these risks. A limitation of this study is the small number of sponges that were evaluated, which may have constrained the levels of sponge microbiome diversity and the number of bacterial foodborne pathogen species observed. Future research should further examine the total and viable microbiomes of kitchen surfaces from a diverse range of households to improve the understanding of indoor surface microbiomes and the risks of viable pathogen exposure and cross-contamination.

## Acknowledgments

The authors thank Brighter Bites staff members and volunteers for their generous assistance. The authors are also grateful to Behzad Imanian and Deena Awad (Institute for Food Safety and Health, Bedford Park, IL) and Diana Stewart (U.S. FDA) for their gracious support during this study. The findings and conclusions in this study are those of the authors and do not necessarily represent the views of the U.S. FDA. This research received no specific grant from any funding agency in the public, commercial, or not-for-profit sectors

All authors contributed to the conception and design of the study. CC collected environmental samples, including dish sponges. CC and JS conducted laboratory experiments. CC performed data analysis. CC and JS created data visualizations. CC wrote the initial draft of the manuscript. All authors contributed manuscript revision and approved the submitted version.

